# Reorganization of thalamocortical connections in congenitally blind humans

**DOI:** 10.1101/449009

**Authors:** Theo F. Marins, Maite Russo, Erika C. Rodrigues, Marina Monteiro, Jorge Moll, Daniel Felix, Julia Bouzas, Helena Arcanjo, Claudia D. Vargas, Fernanda Tovar-Moll

**Affiliations:** D’Or Institute for Research and Education (IDOR), Rio de Janeiro, Brazil; Post-Graduation Program in Morphological Sciences (PCM) of the Institute of Biomedical Sciences (ICB), Federal University of Rio de Janeiro (UFRJ), Rio de Janeiro, Brazil; Institute of Biophysics Carlos Chagas Filho (IBCCF), Federal University of Rio de Janeiro (UFRJ); Centro de Oftalmologia Especializada

**Keywords:** cross-modal plasticity, congenital blindness, diffusion tensor imaging, thalamus, cortical thickness

## Abstract

Cross-modal plasticity in blind individuals has been reported over the past decades showing that non-visual information is carried and processed by ‘visual’ brain structures. However, the structural underpinnings of cross-modal plasticity in congenitally blind individuals remain unclear despite multiple efforts. We mapped thalamocortical connectivity and assessed cortical thickness and integrity of white matter of ten congenitally blind individuals and ten sighted controls. We hypothesized an aberrant thalamocortical pattern of connectivity taking place in the absence of visual stimuli from birth as a potential mechanism of cross-modal plasticity. In addition to the increased cortical thickness of the primary visual cortex and reduced integrity of visual white matter bundles, we observed structural connectivity changes between the thalamus and occipital and temporal cortices. Specifically, the thalamic territory dedicated to connections with the occipital cortex was smaller and displayed weaker connectivity in congenitally blind individuals. In contrast, those connecting with the temporal cortex showed greater volume and increased connectivity compared to sighted controls. The abnormal pattern of thalamocortical connectivity included the lateral and medial geniculate nuclei and the pulvinar nucleus. For the first time in humans, a remapping of structural thalamocortical connections involving both unimodal and multimodal thalamic nuclei has been demonstrated, shedding light on the possible mechanisms of cross-modal plasticity in humans. The present findings may help understand the functional adaptations commonly observed in congenitally blind individuals.

## INTRODUCTION

Blindness and vision impairment affects at least 2.2 billion people around the world^1^. A significant part of these patients has congenital blindness, which can be caused by congenital anomalies and infantile glaucoma, among other causes^2^. Congenital blindness represents an intriguing model for understanding how the brain builds and maintains its fundamental principles of organization and hierarchy of processing in the absence of one of the major sources of input. The impact of congenital blindness on spared sensory systems has also been a focus of study over the past years. A better characterization of the basis of blindness-related brain alterations, including the well-described phenomenon of cross-modal plasticity, can pave the way for developing and optimizing new inclusive devices and brain-machine interfaces.

Numerous brain alterations can occur in response to the lack of visual input, including in white matter (WM) and gray matter (GM) of both cortical and subcortical brain structures^3–5^. Congenitally/early blind individuals often exhibit atrophy of optic chiasm and reduced microstructural integrity of optic radiations and geniculocalcarine tract^6–11^, as investigated by Diffusion Tensor Imaging (DTI). Also, splenium of the corpus callosum, inferior longitudinal fasciculus, and inferior fronto-occipital fasciculus, major WM bundles connecting different cortical areas involved in visual processing and perception, are impacted by the absence of visual input^10,12,–16^. The way congenital and early blindness affect the primary visual cortex structure is a matter of debate. Whereas increased cortical thickness and volume have been reported^17–19^, the opposite effects have also been observed^6,8,20,21^.

Functional neuroimaging studies have shown that visual cortical structures in the blind brain often participate in the processing of non-visual information, which is called cross-modal plasticity. In blind individuals, this reorganization is commonly observed as a visual takeover by the auditory and tactile systems^22–27^. Still, it has also been reported involving language^28,29^, olfaction, and gustation^30,31^ very often leading to increased abilities and behavioral improvement in the remaining senses. In addition, recruitment of ‘visual’ brain areas in blindness has been reported during auditory^18,32,–35^, tactile^23^, and linguistic ^29^ tasks, and correlation with behavioral gains has also been shown^33,36,–39^. Moreover, virtual lesions targeted to the ‘visual’ cortex induced by Transcranial Magnetic Stimulation (TMS) can temporarily impair performance on auditory^12^ and tactile^40^ tasks in blind individuals. Altogether, these findings suggest that the brain undergoes critical changes in the absence of visual input, which pave the way for cross-modal plasticity.

While evidence corroborates cross-modal plasticity in blindness, neuroanatomical correlates underlying this phenomenon are controversial and may involve a wide range of brain structural changes, including the development of new connections and unmasking /rewiring of the existing ones^41^. Indeed, it is unclear how and which pathways convey non-visual information to the visual cortex in the blind brain. Most findings on the possible anatomical underpinnings of cross-modal plasticity and its mechanisms come from animal models of blindness studies. They point to the emergence of direct connections between deprived visual, auditory, and somatosensory cortices, indicating that non-visual information would reach the visual cortex by direct corticocortical connectivity, a transient connection that becomes stabilized during development by the lack of appropriate visual stimuli^42–48^. On the other hand, the preservation of geniculocalcarine pathways in blind individuals^8,49,50^ has been reported, which supports the idea that adaptive plasticity may rely upon the conservation of thalamocortical pathways. In the naturally blind mole rat, the remnant visual pathways carry auditory information to the visual cortex as their visual nuclei of the thalamus receive input from the inferior colliculus, an important auditory center^51,52^ Moreover, in addition to the preserved thalamocortical connections, visual cortical areas receive input from auditory and somatosensory nuclei of the thalamus in blind mice and opossum^46,53^. These findings suggest that robust changes in the thalamocortical connectivity may occur in response to the lack of visual input and could also explain the cross-modal plasticity observed in congenitally blind individuals.

Thus, despite remarkable changes that have been reported in the congenitally blind brain, the structural underpinnings of cross-modal plasticity are still uncertain in humans. In the present study, we combined neuroimaging techniques based on diffusion-weighted imaging, probabilistic tractography, and brain morphometry to interrogate the thalamic (re)mapping of cortical connections of congenitally blind individuals. Specifically, we tested the hypothesis that structural changes of thalamocortical projections involved in visual and multimodal sensory processing would occur in response to the absence of appropriate visual input from birth. In addition, we hypothesized that congenital blindness impairs the appropriate maturation of ‘visual’ areas, reflected in WM and GM changes. Our results point to a remapping of the thalamic connections with both temporal and occipital cortices in congenitally blind individuals compared to a matched sample of sighted controls and an increased cortical thickness in the primary visual cortex (V1) and impaired WM microstructure of visual structures.

## METHODS

### Subjects

This study was conducted in accordance with the ethical standards-compliant of the Declaration of Helsinki and has been approved by the IDOR / Copa D’Or Ethics and Scientific Committee. Ten congenitally blind (CB; mean age: 31.8, standard deviation: 8.7, 6 males) individuals and ten sex- and age-matched sighted controls (SC; mean age: 32.2, standard deviation: 6.66, 6 males) were included in the study. All participants were right-handed, had no history of neurologic or psychiatric diseases, and were not taking brain-active medication. For the inclusion of the CB individuals, their medical records and additional clinical exams (such as the visual evoked potentials) have been reviewed. Afterward, they were examined by an experienced ophthalmologist, confirming that they were totally blind or had only minimal residual light sensitivity. All blind participants were Braille readers. The cohort characteristics are summarized in Table 1.

**Table 1.** Cohort characteristics

| Patient ID | Age | Gender | Cause of congenital blindness | Handedness | Braille |
| --- | --- | --- | --- | --- | --- |
| CB01 | 35 | F | Anophthalmia | Right | Yes |
| CB02 | 36 | M | Macular dystrophy | Right | Yes |
| CB03 | 32 | F | Glaucoma | Right | Yes |
| CB04 | 18 | M | Glaucoma | Right | Yes |
| CB05 | 40 | M | Glaucoma | Right | Yes |
| CB06 | 27 | M | Glaucoma | Right | Yes |
| CB07 | 22 | M | Retinal detachment | Right | Yes |
| CB08 | 44 | M | Unknown | Right | Yes |
| CB09 | 24 | F | Retinopathy of prematurity | Right | Yes |
| CB10 | 40 | F | Unknown | Right | Yes |

M: male. F: female

### Data Acquisition

Brain imaging was performed at D’Or Institute for Research and Education (IDOR) in a 3T Achieva scanner (Philips Medical Systems, the Netherlands) using an eight-channel SENSE head coil. Imaging consisted in a high-resolution 3D T1-weighted image (1mm^3^ isotropic, TR/TE (ms) = 7.2 / 3.4, FOV= 240 x 240, matrix = 240, slice thickness = 1mm) and diffusion-weighted (2 x 2 x 2mm^3^ isotropic, no gap, TR/TE (ms)= 10150/60, FOV = 224 x 224, matrix = 112 x 112) with diffusion sensitization gradients applied in 64 noncollinear directions, with a b factor of 1000 s/mm^2^, one volume without diffusion weighting.

### Data Analysis

Before data preprocessing, images were visually inspected for excessive movements or artifacts. Diffusion-weighted images were preprocessed and analyzed using FSL^54^ toolboxes. In each subject, original data were corrected for the effects of head movement and eddy currents using eddy correct, and a brain mask was created by running BET^55^ on the B=0 (no diffusion weighting) image. We created FA images using FDT (with DTIFIT algorithm), part of FSL^56^. All subjects’ data were aligned into standard space (Montreal Neurological Institute, MNI) using the nonlinear registration tool FNIRT, and the mean FA image was created and thinned to create a mean FA skeleton. The FA map of each subject was projected onto skeletonized FA, and the resulting data was fed into voxel-wise cross-subject statistics using TBSS^57^. We tested for group differences with an unpaired t-test using Randomise^58^ for permutation-based (5000 permutations) non-parametric testing of whole skeleton FA. Also, analysis restricted to the whole thalamus was separately conducted.

Connectivity-based segmentation of the thalamus was also performed using FDT, as described previously^59^, in each hemisphere separately. Six cortical masks were predefined using the Harvard-Oxford atlas: Prefrontal, Precentral, Postcentral, Posterior Parietal (parietal cortex, except for the postcentral cortex), Occipital, and Temporal. Since the Oxford Thalamic Connectivity Probability Atlas does not include the lateral and medial geniculate nuclei (LGN and MGN, respectively), both right and left thalamic masks were obtained from an atlas on the Colin-27 Average Brain^60^. All masks were registered into the subjects’ space using nonlinear registration. After BEDPOSTX^56^, probabilistic tracking was conducted with the PROBTRACKX^56^ tool, in which the unilateral thalamus was defined as the seed region and the six ipsilateral cortical masks as classification targets. Following, we calculated the number of samples reaching each target mask as a proportion of the total number of samples reaching any target mask. Then, the hard segmentation of each thalamus based on its connectivity to each of the six ipsilateral cortical areas was performed. The volume of each resultant segment of the thalamus was normalized by the volume of the ipsilateral thalamus **at** the subject level. Group analysis of the normalized volume was performed with SPSS 20.0 (IBM Corporation, New York) using a repeated-measures analysis of variance (ANOVA; within-subjects factors: ‘segment’ and ‘side’; between-subjects factor: ‘group’). We used Bonferroni to adjust for multiple comparisons.

To perform a voxel-wise analysis of the thalamocortical connectivity and compare them between groups, the resultant six segments at the subject level were transformed back to the MNI space. Each segment was overlapped to create a single mask, thus consisting in a sum of all individual segment masks from both groups. The voxel-wise analysis was conducted within this mask separately for each thalamic segment. Group differences were investigated with an unpaired t-test using Randomise^58^ for permutation-based (5000 permutations) non-parametric testing of the connectivity maps for each segment.

### Cortical thickness analysis

Cortical thickness analysis was performed based on high-resolution 3D T1-weighted gradient-echo images using Freesurfer software (http://surfer.nmr.mgh.harvard.edu/). Preprocessing steps included removal of non-brain tissue, segmentation of the subcortical white matter and deep gray matter volumetric structures, intensity normalization, tessellation of the gray matter/white matter boundary, automated topology correction, and surface deformation following intensity gradients.

To investigate group differences, we applied the command-line Freesurfer group analysis stream, including correction for multiple comparisons. First, the data from every subject was assembled by resampling each subject’s data onto a common space, spatially smoothing it, and concatenating all the subject’s resampled and smoothed data into a single file. We defined the design matrix and design contrast comparing both groups, resulting in a t-test analysis of the whole brain. The statistical results were then submitted to a cluster-wise correction for multiple comparisons (p<0.05).

For simplicity, group differences are referred to as “increase” and “decrease” throughout the manuscript taking the SC group as a reference. The coordinates in the figures are given according to MNI space, and results are plotted on the MNI standard brain.

### Data availability

Data that support the present findings are available from the corresponding author upon reasonable request.

## RESULTS

### Reorganization of thalamocortical connections

The thalamus segmentation based on the structural connectivity patterns to six predefined cortical areas (Prefrontal, Precentral, Postcentral, Posterior Parietal, Temporal, and Occipital) successfully resulted in six thalamic segments on each side (Fig. 1) in all participants.

**Fig. 1.**
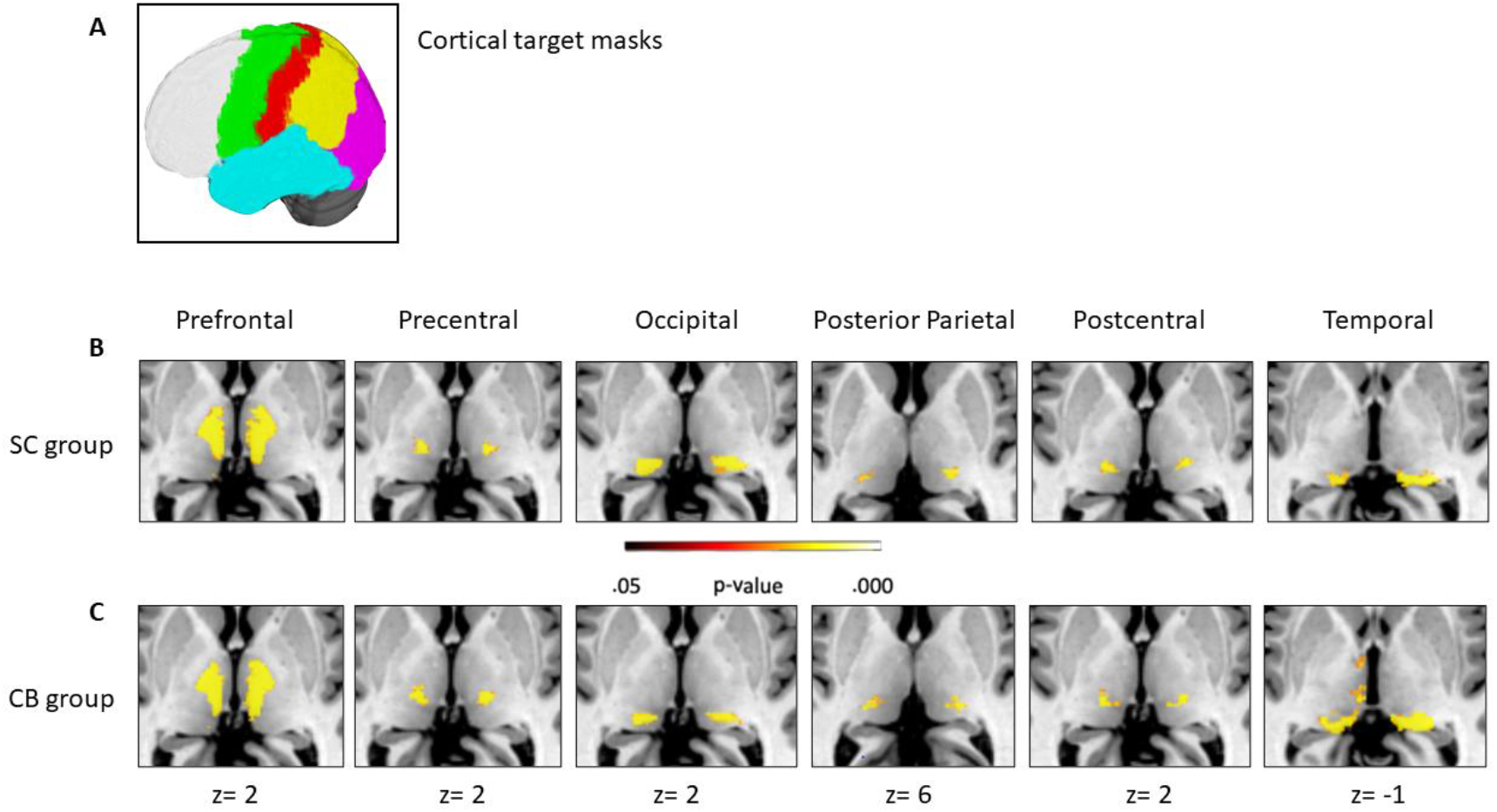
Thalamocortical Connectivity in Sighted Controls and Congenitally Blind Individuals. (A) Six cortical masks were predefined using the Harvard-Oxford atlas: Prefrontal, Precentral, Postcentral, Posterior Parietal (parietal cortex, except for the postcentral cortex), Occipital, and Temporal. Resultant thalamic segments showing statistically significant connectivity to each cortical area in the (B) SC (sighted controls, top) and (C) CB (congenitally blind, middle) groups. The color bar represents the p-value after FWE correction. The normalized volume of the Occipital and Temporal segments showed group differences (p<0.001, Bonferroni correction for multiple comparisons).

The analysis of thalamic volumes revealed an interaction effect between ‘segment*group’ (F(2.32, 41.80)=4.76, p=0.01). *Posthoc* analysis with Bonferroni correction for multiple comparisons on the interaction ‘segment*group’ indicated that the SC and CB groups significantly differed in the volume of the Occipital segment (p=0.013), being reduced in the CB group; and Temporal segment (p=0.002) being increased in the CB group (Fig. 1; Table 2).

**Table 2.** Normalized volume comparisons of thalamic segments

| Segment | Side | SC |  | CB |  |
| --- | --- | --- | --- | --- | --- |
|  |  | Mean<br>Volume | SD | Mean<br>Volume | SD |
| Prefrontal<br>p=0.064 | R | 0.561224 | 0.0372437 | 0.515953 | 0.0578148 |
|  | L | 0.537903 | 0.0424005 | 0.505637 | 0.0457513 |
| Precentral<br>p=0.14 | R | 0.055635 | 0.0125891 | 0.060761 | 0.0150288 |
|  | L | 0.067851 | 0.0290276 | 0.078154 | 0.0259177 |
| Occipital*<br>p=0.013 | R | 0.11052 | 0.0286398 | 0.071793 | 0.0451293 |
|  | L | 0.138718 | 0.0443775 | 0.088397 | 0.0174996 |
| Posterior Parietal<br>p=0.97 | R | 0.065566 | 0.0267344 | 0.064867 | 0.0174419 |
|  | L | 0.049694 | 0.026557 | 0.05505 | 0.0229696 |
| Postcentral<br>p=0.07 | R | 0.044252 | 0.0156764 | 0.040539 | 0.0148974 |
|  | L | 0.060081 | 0.0209732 | 0.071497 | 0.0407492 |
| Temporal*<br>p=0.002 | R | 0.162803 | 0.0458977 | 0.246086 | 0.0842971 |
|  | L | 0.145753 | 0.0363323 | 0.201264 | 0.0570562 |

\*Segments that showed significant ( $p < 0.05$ ) between-group differences after Bonferroni correction for multiple comparisons.
L= left thalamus; R= right thalamus. Mean volume refers to the normalized mean volume in arbitrary units.

In addition, there was a main effect of ‘segment’ (F(2.32, 41.80)=635,92, p<0.005) and an interaction between ‘segment*side’ (F(2.56 46.01)=9.77, p<0.005). Pairwise comparisons were run to investigate the main effect of ‘segment’ and interaction ‘segment*side’ adjusting for Bonferroni (Table S1 and S2).

To investigate whether congenital blindness alters the anatomical connectivity pattern between the thalamus and the cortex, we conducted a group comparison using threshold-free cluster enhancement (TFCE, FWE-corrected, *p*<0.05) of the probability maps of connectivity of each thalamic segment. This analysis revealed that the connectivity of the thalamic segment connecting with the occipital and the temporal cortices significantly differed between groups, with the thalamo-temporal connectivity being increased and the thalamo-occipital connectivity being reduced in the CB group. Specific regions showing increased thalamo-temporal connectivity in the CB group were observed in both thalami, specifically in the bilateral LGN, bilateral pulvinar, bilateral MGN, left medial dorsal nucleus, left anterior nucleus, left ventral anterior nucleus and left lateral posterior nucleus (Fig. 2). On the other hand, reduced thalamo-occipital anatomical connectivity in the CB group was restricted to the territory of the pulvinar and lateral posterior nucleus on the left thalamus (Fig. 2). Further analysis revealed that both group differences in thalamocortical connectivity (increased thalamo-temporal connectivity and decreased thalamo-occipital connectivity) partially shared a common neural territory on the left pulvinar (Fig. 2). Thalamic projections to the Prefrontal, Precentral, Postcentral, and Posterior Parietal segments did not show statistically significant differences between groups.

**Fig. 2.**
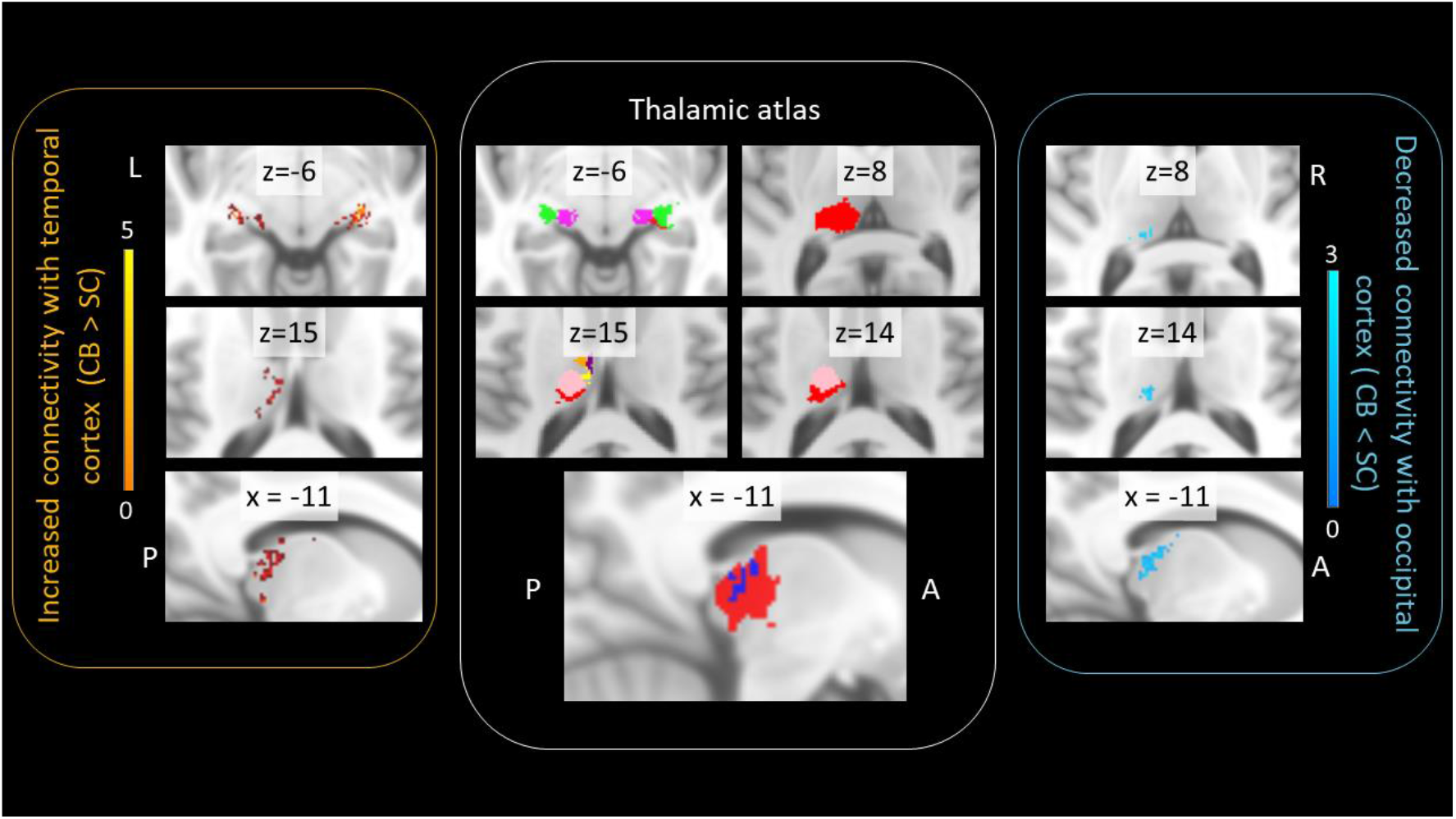
Thalamocortical Connectivity Changes in Congenitally Blind Individuals. The yellow box (left) depicts thalamic areas that exhibited increased connectivity with the temporal cortex, including MGN, LGN and pulvinar bilaterally. The blue box (right) depicts thalamic territory that exhibited decreased connectivity with the occipital cortex in congenitally blind individuals, namely the left pulvinar/lateral posterior nucleus. The white box (middle) shows thalamic territories obtained from an atlas based on the Colin-27 Average Brain^88^ and depicts the location of LGN (green), MGN (dark pink), pulvinar (red), medial dorsal (yellow), ventral anterior (orange), anterior (purple), and lateral posterior (light pink) nuclei. A graphical overlay (dark blue) of thalamic areas that exhibited both increased connectivity to the temporal cortex and decreased connectivity to the occipital cortex (p<0.05, FWE-corrected) in CB individuals is shown (white box, bottom). L = left; R = right; A = anterior; P = posterior. The coordinates are given according to the MNI space and plotted on the MNI standard brain. Color bars represent the t-value.

### White Matter Microstructure Changes in Blind Subjects

We investigated differences in WM microstructure using TBSS for fractional anisotropy (FA). Whole-brain (threshold-free cluster enhancement (TFCE), corrected *p*<0.05) analysis showed decreased FA in the major WM structures connecting the occipital cortex, such as the splenium of the corpus callosum, bilateral inferior longitudinal fasciculi, bilateral inferior fronto-occipital fasciculi, and bilateral superior longitudinal fasciculi (Fig. 3) in the CB group. Additional analysis restricted to the whole thalamus revealed a diffuse decrease in FA in the CB group, including left and right pulvinar, medial dorsal nucleus, right ventral lateral nucleus, right lateral posterior, and left lateral nucleus (Fig. 3).

**Fig. 3.**
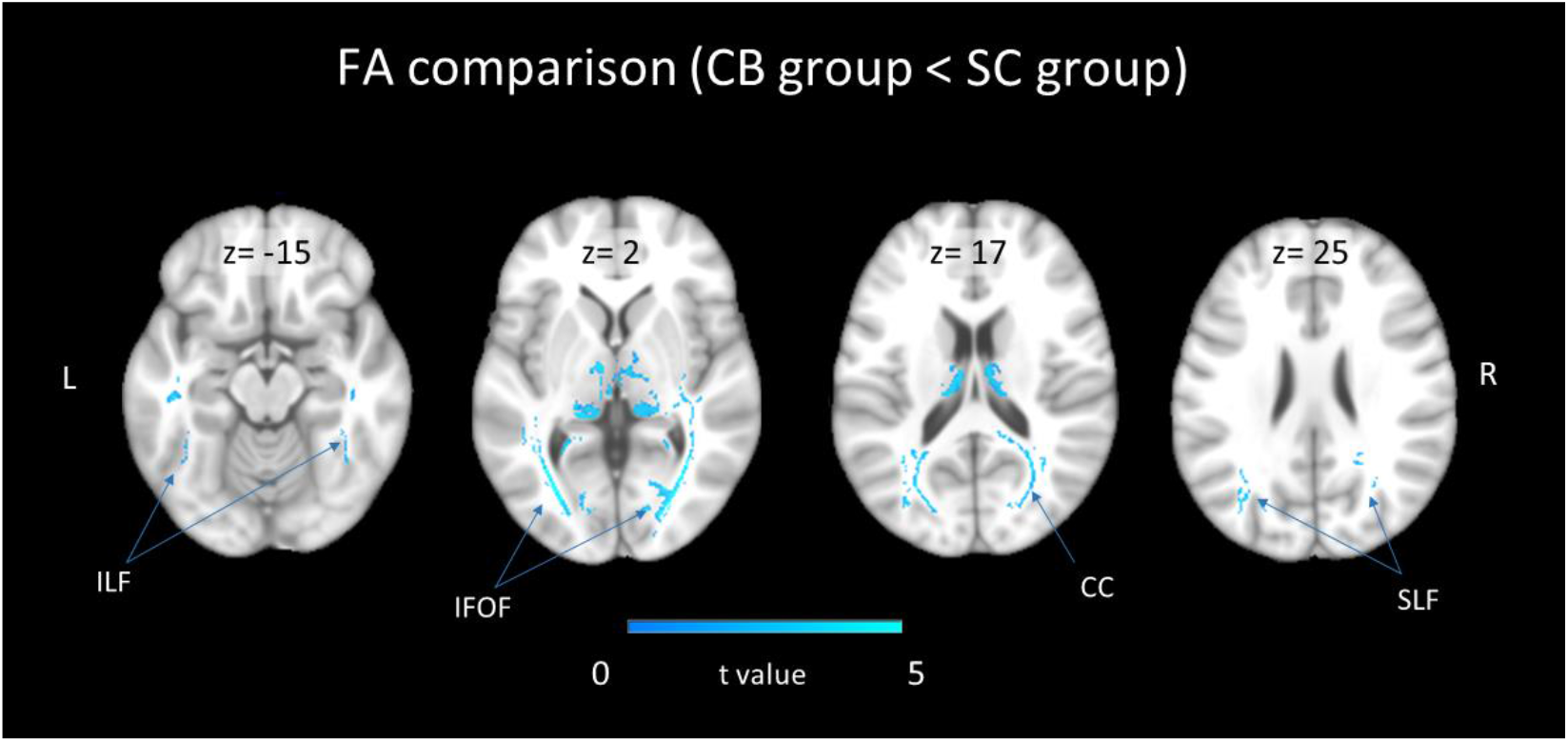
FA comparison (CB group < SC group). White matter tracts (whole brain) showed reduced FA (fractional anisotropy) in the CB (congenitally blind) group, as compared to SC (sighted controls) group FWE-corrected, p< 0.05). FA difference in the thalamus, resulting from a separate analysis, is also displayed. The color bar represents the t value corrected for multiple comparisons. ILF = inferior longitudinal fasciculus; IFOF = inferior fronto-occipital fasciculus; CC = corpus callosum; SLF = superior longitudinal fasciculus.

### Increased Cortical Thickness in Blind Subjects

Whole-brain comparison of cortical thickness (whole-brain analysis, cluster-corrected for multiple comparisons, *p* < 0.05; Fig. 4A) revealed greater thickness of the left primary visual cortex (V1, Brodmann area 17 (BA17), Size= 268.24 mm^2^, X=-5.2, Y=-86.4, Z=10.3, MNI) in the CB group (CB group > SC group). This single cluster was then masked to further investigate thickness in each participant (Fig. 4B). No other cortical region showed group differences in thickness.

**Figure 4.**
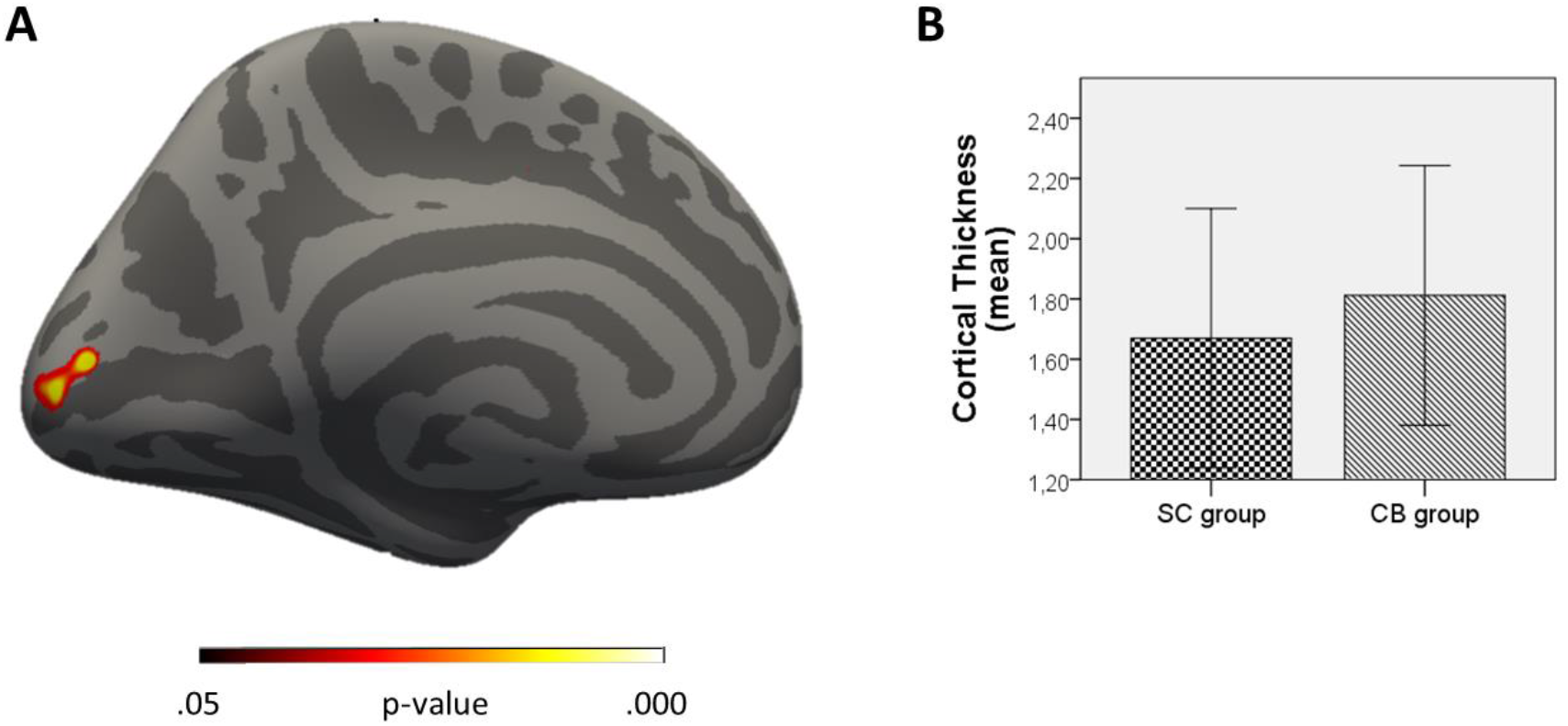
Increased Cortical Thickness in the Congenitally Blind Group. Increased cortical thickness in the left V1 (BA17) in the CB (congenitally blind) group compared to the SC (sighted controls) group, whole-brain analysis. (B) The mean cortical thickness (in mm) of each group is displayed for visualization of the clustered values and not for inference (error bar represents standard deviation).

## DISCUSSION

The present findings indicate that the absence of appropriate visual input from birth in humans leads to a wide range of structural brain changes. For the first time, we described a structural remapping of thalamocortical connectivity, which may help understand the functional adaptations commonly observed in congenitally blind individuals.

The analysis of a sample of right-handed CB individuals, Braille readers, brought new evidence of reduced thalamic areas projecting to/from the occipital cortex compared to SC. On the other hand, a greater volume of the thalamic territory dedicated to connections with the temporal cortex was observed in CB compared to SC. Additionally, the voxel-wise analysis revealed that thalamic nuclei such as bilateral LGN, MGN, and pulvinar were more connected to the temporal cortex in CB. In contrast, left pulvinar and lateral posterior nuclei displayed weaker projections to the left occipital cortex in this group. Interestingly, we found strengthened connectivity of the left pulvinar to the temporal cortex over the occipital cortex, suggesting a possible rerouting of the pulvinar connections in CB. Moreover, we confirmed previous findings by showing that visual cortical areas exhibit greater thickness, besides a diffuse impairment in the microstructure of major visual WM bundles.

Our findings bring new evidence in line with previous studies showing impairment of thalamic nuclei involved in visual and multimodal processing, such as LGN and pulvinar. Using voxel-based morphometry (VBM), previous studies have shown volume reductions in thalamic visual centers in CB individuals, including dorsal LGN and pulvinar^13,61^, identified with the aid of brain atlases. A similar approach has been used to describe volume reduction in WM close to the LGN in six bilaterally anophthalmic subjects compared with sighted subjects^8^. However, it is worth noting that the present study was not limited to comparing the volume of thalamic structures between groups. Instead, we went beyond by mapping the thalamocortical projections and exploring a possible remapping of thalamic territories and their connectivity pattern to/from the cortex. Thus, one key difference between the present and previous studies is that visual-related thalamic nuclei investigated here were identified using subject-level tractography. Interestingly, the thalamic territory occupied by projections to/from the temporal cortex showed greater volume, whereas those to/from the occipital cortex showed reduced volume in CB individuals compared to the SC, which brings new insights into the thalamic remodeling in congenital blindness. The comparison of thalamic volumes obtained from different approaches, such as probabilistic tractography and VBM, must be explored in future studies considering their technical aspects and biological underpinnings. For example, we employed a probabilistic tractography algorithm to map the connectivity distributions from individual voxels within the thalamus in each participant, while VBM relies on the boundaries between thalamic nuclei^62^.

Even though interesting, volume comparisons of thalamic structures have low intrinsic sensitivity, as differences in connectivity may not be reflected as volume changes. This would be even more tricky since thalamocortical remapping may be presented as both shrinkage and expansions of nuclei territories in a non-homogeneous and distributed pattern. For the first time, we disentangled this by comparing the voxel-wise differences in thalamocortical connectivity between groups, revealing the pulvinar’s central role in remapping thalamocortical connectivity. WM connections between pulvinar/lateral posterior nuclei and occipital cortex were decreased in the CB group. On the other hand, increased connectivity to/from the temporal cortex was observed in the pulvinar. These findings suggest a thalamic shift of cortical connections in blind individuals, prioritizing the temporal projections over the occipital ones.

The pulvinar is the largest, higher-order multimodal thalamic nucleus, involved mainly in visual attention^63^. It displays a wide range of cortical connectivity beyond the visual cortex, roughly to the gray matter of all lobes and subcortical structures such as the amygdala and superior colliculus^63–65^. Evidence suggests that cortico-cortical integration via pulvinar is an important pathway for transferring visual information between cortical areas^65,66^. Thus, it is reasonable to hypothesize that the pulvinar may become a central player in the dynamics of brain plasticity, possibly by integrating different sensory modalities at associative levels and paving multimodal plasticity in blind individuals. However, the pulvinar typically connects to higher-order areas of the temporal cortex such as inferior, ventral, and lateral temporal areas, temporoparietal junction, and mesial temporal cortex^64,67,68^. Therefore, it is impossible to conclude whether the aberrant connectivity pattern of the pulvinar observed here relies on new thalamocortical projections to the temporal cortex or a mechanism of remodeling the preexisting ones. Hence, the role of pulvinar in cross-modal plasticity needs to be considered and further explored in future studies.

It is worth noting that our findings add new insights to the still-developing literature on thalamocortical connectivity changes in blind humans. Even though Reislev and colleagues^69^ have investigated possible differences in thalamocortical connectivity between CB and SC groups, evidence of thalamic remapping has not been found. For the first time using a voxel-wise approach, we reported converging evidence that thalamic projections to/from the temporal cortex are increased as opposed to those to/from the occipital cortex. First, comparisons of the thalamic territory connecting to the temporal and occipital cortices showed increased and decreased volume in the CB group, respectively. Second, the voxel-wise analysis pointed to an aberrant pattern of thalamocortical connectivity involving pulvinar/lateral posterior nucleus, LGN, and MGN, among others. Thus, despite similarities, we used original analysis to describe the impact of congenital blindness on the thalamocortical connectivity in humans.

In the past years, several authors have described increased V1 thickness in the blind brain^18,19,39,70^, which can be explained by the lack of correct pruning during ontogenesis due to insufficient input^19,71,72^. On the other hand, the absence of appropriate input may also result in an impaired structure of V1^6,8,15,21^. In the present study, the CB group displayed greater thickness of the left but not the right V1 (BA 17) than SC. Although the changes of cortical thickness in blind individuals have been reported bilaterally in previous studies, the most prominent results are often reported in the left occipital cortex in right-handers^18,70^, indicating lateralization effects of plasticity in these cases. Indeed, the visual cortex is highly lateralized for a wide range of stimuli^73–76^. Non-visual tasks, such as spatial and pitch discrimination of sounds^34^, voice perception^77^ and recognition^35^, sound localization^36^, syntactic^29^, and semantic^28^ processing of sentences, elicit lateralized visual cortex activation in blind individuals. In addition, the lateralization of functional connectivity to associative brain areas has also been reported^34,78^. Due to the nature of hemispheric dominance of the visual cortex that differently affects both hemispheres depending on the task, it is hard to define precisely what aspects of brain plasticity underpin the left lateralization of increased cortical thickness in CB. One possible mechanism affecting the left V1 is that input to V1 seems to be a critical part of cortical maturation and pruning^19,71,72^. Indeed, we found evidence of left-lateralized impairment effects on thalamo-occipital connections in CB. Thus, the involvement of thalamocortical connectivity in this phenomenon should be addressed in future studies.

We did not find evidence of thalamic remapping of connections to the somatosensory and motor cortices, for example, in CB. However, careful interpretation of the results must consider two key points. First, the connectivity-based segmentation of the thalamus used in the present study is a ‘winner-takes-all’ approach in which the most probable connections are considered while the least probable ones are discarded. Hence, plasticity involving somatosensory and motor remodeling of thalamic connections may not have been strong enough to be detected by our methods. As a result, the ‘normal’ pattern of thalamic segmentation would mask meaningful but less structured connections. For example, even though weaker, aberrant somatosensory, auditory and motor thalamic nuclei are connected with V1 in blind opossum^46^. In addition, V1 activation following tactile stimulation may be driven by aberrant connections between the ventroposteriorlateral nucleus and LGN in CB individuals^79^. Second, although we did not intend to assess corticocortical connections between visual and the remaining sensory modalities, this kind of direct connectivity has been reported in animal models of blindness and must be considered a possible mechanism of cross-modal plasticity in humans. In fact, the intermodal connections during ontogenesis seem to be altered by the lack of visual input in animals^48,53,80–82^. In humans, indirect measurement based on functional connectivity pointed to direct connections between the occipital and auditory cortex^83^ and primary somatosensory cortex^84,85^. Moreover, the visual cortex of enucleated animals also displays both abnormal corticocortical and thalamocortical connections with somatosensory, auditory, and motor areas^46^, suggesting that both mechanisms may coexist to support cross-modal plasticity in the blind brain.

In accordance with previous studies that showed decreased FA of WM in CB^10,13–15^, we observed focal impairment of WM integrity in CB individuals impacting most occipital connections to the associative cortical areas. Structures showing decreased FA were splenium of the corpus callosum, bilateral inferior longitudinal fasciculi, bilateral inferior fronto-occipital fasciculi, and bilateral superior longitudinal fasciculus, which are known for their role in processing and integration of visual information. Lower FA values refer to WM integrity and myelination impairments and are commonly pointed to as a biomarker of developmental changes, axonal degeneration, and plasticity. Indeed, visual input is crucial for maturation and refinement of early crude connectivity of the visual system,^71,82,86^ suggesting that decreased myelination of visual WM tracts in CB individuals might reflect the lack of maturation of the system rather than axonal degeneration per se. Moreover, the thalamus also displayed reduced integrity in the CB group, indicating that microstructural impairment is not restricted to corticocortical bundles. FA analysis of GM structures may represent a measure of cytoarchitecture^87^ and is in line with previous studies^69^. In addition to the reduced FA, previous studies suggest that the lack of visual input impacts WM volume compared with sighted controls^7,8,16^.

The present findings raise intriguing questions that should be addressed in future studies. For example, the CB group showed increased structural connectivity between the ventral anterior nuclei and the temporal cortex. Given that most studies focused on exploring abnormal thalamocortical connectivity involving V1, the possible involvement of a motor thalamic nucleus, such as the ventral anterior, deserves further investigation. In addition, our sample of blind individuals includes only CB. Despite similar approaches have targeted both CB and late blind individuals^69^, future studies should employ voxel-wise investigation of thalamocortical connectivity in people with acquired blindness to elucidate to what extent the present thalamocortical connectivity changes depend on visual experience.

In summary, the present study corroborates previous findings pointing to the blind brain as a model of a wide range of plastic phenomena. For the first time in humans, the remapping of thalamocortical connections involving both unimodal and multimodal thalamic nuclei is described, which may represent a mechanism of how non-visual stimuli are relayed to the ‘visual’ cortex. Future studies should employ neurophysiologic approaches to correlate these changes with functional plasticity often observed in the absence of visual stimuli from birth.

## Abbreviations

WM: White Matter
DTI: Diffusion Tensor Imaging
LGN: Lateral Geniculate Nucleus
MGN: Medial Geniculate Nucleus
CB: Congenitally blind
SC: Sighted controls

## ACKNOWLEDGEMENTS

We thank Ivanei Bramati and Débora Lima for their support during data analysis and writing. The authors declare that the research was conducted in the absence of any commercial or financial relationships that could be construed as a potential conflict of interest.

## Supplementary material

**Table S1.** Investigation of the effect of ‘segment’.

| Segment (pairwise comparison) |  | p-value (adjusted) |
| --- | --- | --- |
| Frontal<br>mean = 0.539 (sd = 0.049) | Motor | $4.55 \times 10^{-130*}$ |
| | Posterior Parietal | $3.2 \times 10^{-132*}$ |
| | Somatosensory | $3.01 \times 10^{-133*}$ |
| | Occipital | $1.49 \times 10^{-122*}$ |
| Motor<br>mean = 0.064 (sd = 0.024) | Occipital | $7.36 \times 10^{-4*}$ |
|  | Posterior Parietal | 1 |
|  | Somatosensory | 1 |
| | Temporal | $2.2 \times 10^{-32*}$ |
| Occipital<br>mean = 0.1 (sd = 0.039) | Posterior Parietal | $4.3 \times 10^{-4*}$ |
| | Somatosensory | $1.28 \times 10^{-5*}$ |
| | Temporal | $6.62 \times 10^{-18*}$ |
| Posterior Parietal<br>mean = 0.058 (sd = 0.024) | Somatosensory | 1 |
| | Temporal | $1.39 \times 10^{-33*}$ |
| Somatosensory<br>mean = 0.047 (sd = 0.021) | Temporal | $1.65 \times 10^{-37*}$ |
| Temporal<br>mean = 0.193 (sd = 0.063) | Frontal | $4.04 \times 10^{-101*}$ |

Mean and standard deviation (sd) refer to the normalized mean volume in arbitrary units. P values were adjusted for Bonferroni correction. Asterisks indicate statistically significant differences.

**Table S2.** Investigation of the interaction ‘segment*side’.

| **Segment** | **Volume (mean(sd))** |  | **p-value**<br>**(adjusted)** |
| --- | --- | --- | --- |
|  | **Left side** | **Right side** |  |
| Frontal | 0.536 (0.053) | 0.541 (0.046) | 0.766 |
| Motor | 0.074 (0.027) | 0.053 (0.014) | 0.00308* |
| Occipital | 0.11 (0.035) | 0.09 (0.041) | 0.112 |
| Posterior Parietal | 0.051 (0.025) | 0.064 (0.021) | 0.0812 |
| Somatosensory | 0.059 (0.021) | 0.036 (0.012) | 0.000101* |
| Temporal | 0.169 (0.056) | 0.216 (0.063) | 0.0171* |

Volume and standard deviation (sd) refer to the normalized mean volume in arbitrary units. P values were adjusted for Bonferroni correction. Asterisks indicate statistically significant differences.

